# Computational Fontan Analysis: Preserving accuracy while expediting workflow

**DOI:** 10.1101/2021.10.28.466272

**Authors:** Xiaolong Liu, Seda Aslan, Byeol Kim, Linnea Warburton, Derrick Jackson, Abir Muhuri, Akshay Subramanian, Paige Mass, Vincent Cleveland, Yue-Hin Loke, Narutoshi Hibino, Laura Olivieri, Axel Krieger

## Abstract

**Background:** Post-operative outcomes of the Fontan operation have been linked to graft shape after implantation. Computational fluid dynamics (CFD) simulations are used to explore different surgical options. The objective of this study is to perform a systematic *in vitro* validation for investigating the accuracy and efficiency of CFD simulation to predict Fontan hemodynamics.

**Methods:** CFD simulations were performed to measure indexed power loss (iPL) and hepatic flow distribution (HFD) in 10 patient-specific Fontan models, with varying mesh and numerical solvers. The results were compared with a novel *in vitro* flow loop setup with 3D printed Fontan models. A high-resolution differential pressure sensor was used to measure the pressure drop for validating iPL predictions. Microparticles with particle filtering system were used to measure HFD. The computational time was measured for a representative Fontan model with different mesh sizes and numerical solvers.

**Results:** When compared to *in vitro* setup, variations in CFD mesh sizes had significant effect on HFD (p = 0.0002) but no significant impact on iPL (p = 0.069). Numerical solvers had no significant impact in both iPL (p = 0.50) and HFD (P = 0.55). A transient solver with 0.5 mm mesh size requires computational time 100 times more than a steady solver with 2.5 mm mesh size to generate similar results.

**Conclusions:** The predictive value of CFD for Fontan planning can be validated against an *in vitro* flow loop. The prediction accuracy can be affected by the mesh size, model shape complexity and flow competition.

## Introduction

Fontan surgery is performed as the final step in surgical palliation of single ventricle heart disease. By surgically connecting the inferior vena cava to the pulmonary arteries and bypassing the heart, the pulmonary and systemic circulations are separated, generally allowing for more efficient oxygen delivery for the patient. The shape and implantation of Fontan grafts affects long-term Fontan outcomes (1–3), including pulmonary arteriovenous malformations (PAVM) (2), decreased exercise capacity, underdeveloped pulmonary arteries, and thrombosis. These complications are linked to hemodynamics in post-operative Fontan geometries (3). The correlation between the development of PAVM and hepatic flow distribution (HFD) (4–6) and the negative correlation between decreased exercise capacity and power loss in Fontan (7) have been demonstrated by various groups.

Predicting the best shape and implantation of the Fontan graft involves balancing several different, sometimes opposing goals, including but not limited to balancing HFD and reducing power loss (8, 9). Computational fluid dynamics (CFD) simulation has been employed to explore different surgical options and to assist in predicting post-operative hemodynamic performance (10, 11). Previously, the reliability of CFD simulations has been linked mainly to the solver assumptions and boundary conditions (BC) (12). Implantation of the Fontan grafts as prescribed by the pre-operative surgical plan relies on the surgeon’s experience (13). In addition, even if perfect Fontan graft implantation can be achieved, the geometry of the vessel will be deformed due to the growth of the patient.

Realistic use of preoperative Fontan planning requires development of a tool that is quick, easy to use, requires standard clinical data for its inputs, and can turnaround accurate recommendations quickly and clearly to ensure accurate implantation of Fontan designs. Prior research demonstrated discrepancies between the pre-operative and post-operative results (6), however, it is difficult to separately quantify the relative contribution of errors in CFD prediction, surgical implantation, and vessel structure alternation to the difference between predicted and observed hemodynamics. Additionally, there is a “law of diminishing returns” when using additional and more powerful computational resources which require time and expense for surgical planning. Therefore, it is crucial to explicitly analyze each error source for improving the surgical planning accuracy.

The reliability of CFD simulations is the foundational step towards accurate Fontan surgical planning. As the accuracy of CFD simulations is a primary concern in the U.S. Food and Drug Administration (FDA) regulatory practice, experimental validations of CFD have been conducted in many research studies to characterize the flow field. Computational imaging techniques and/or optical imaging methods (14) are used to visualize *in vivo* and *in vitro* flow field. 4D flow MRI data have been applied to validate hemodynamic parameters computed from CFD simulation such as flow velocities (15), hepatic flow distributions (16–18), wall shear stress (19), viscous dissipation (20).

The objective of this study was to investigate the accuracy and efficiency of CFD simulation to predict Fontan hemodynamics using an *in vitro* mock circulatory benchtop setup. The impact of mesh size and numerical solver on model accuracy were evaluated to understand overall computational efficiency. In addition, this study provided a template to validate CFD simulation with similar considerations in other cardiovascular surgical planning applications beyond Fontan surgery.

## Methods

### Data Acquisition, Image Segmentation, and 3D Model Reconstruction

With the approval of the Institutional Review Board (IRB), anonymized data from MRIs of 10 patients with Fontan circulation were used to create 3D anatomic replicas of the Fontan and proximal thoracic vasculature. Specifically, angiography data with late-phase, non-gated, and breath-held acquisition with resolution 1.4 × 1.4 mm were used to segment and create the 3D replica using medical image segmentation software, Mimics (Materialise, Leuven, Belgium) and our lab’s standard approach (8) (Supplementary material Figure S1). Phase contrast images with resolution 1.5 – 2.2 mm, velocity encoding ranging 100-150 cm/s and 30 reconstructed phases were used to extract flow curves for inlet and outlet BC specific to the patient. Attributes of the 10 patient-specific models are illustrated in Figure 1. The BC obtained from the patient’s *in vivo* 4D flow MRI data were directly used in the Fontan CFD analysis.

**Figure 1.**
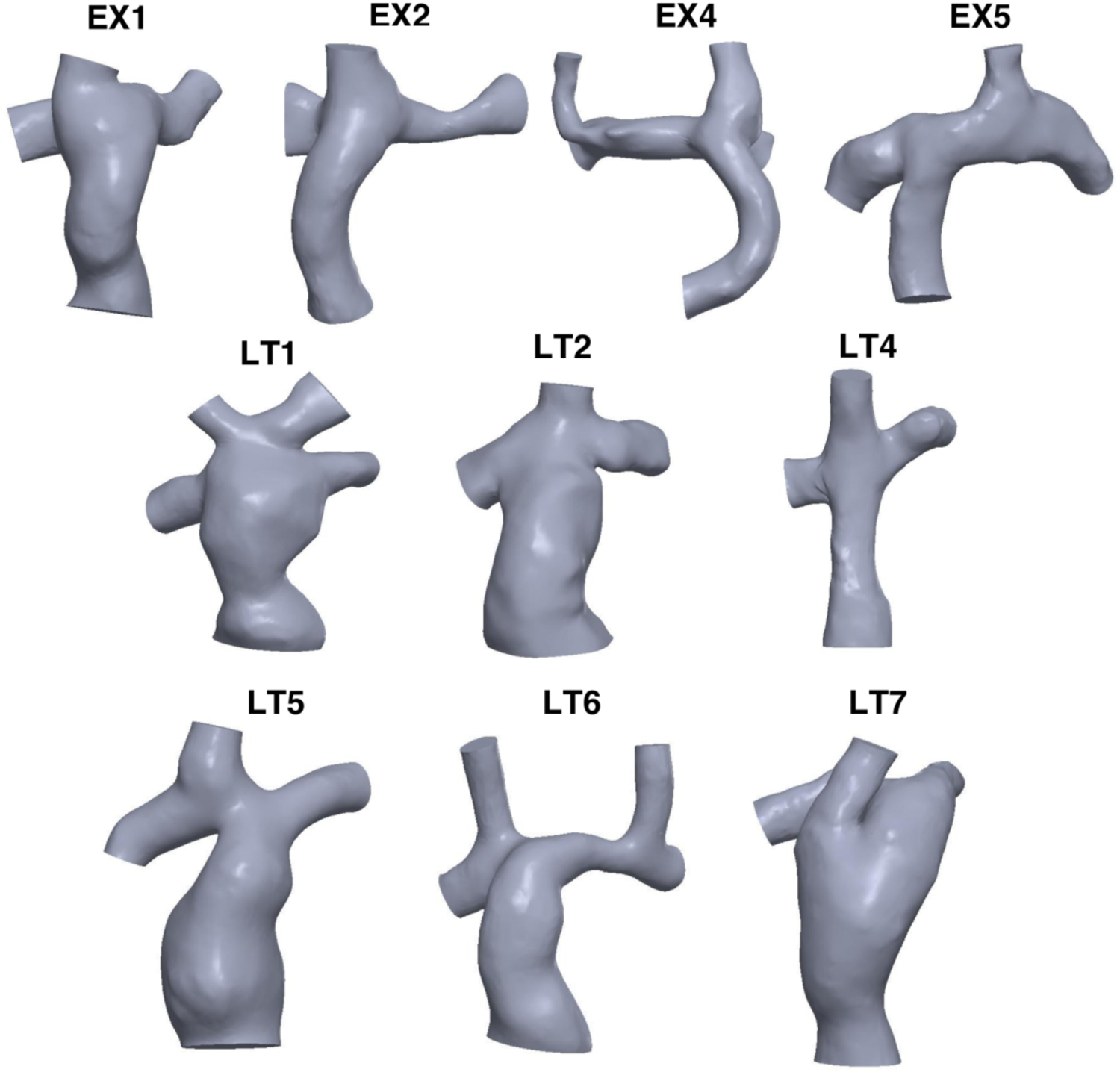
The Cohort of 3D reconstructed Fontan models. EX and LT represent extracardiac and lateral tunnel, respectively. All the models were pre-precessed by making clean cuts at IVC, SVC, LPA and RPA for defining the areas of inlet/outlet BC. The cohort represents a variety of model geometry, different numbers of inlets, and different anatomy conditions, such as the existence of coartation in EX2 and EX5.

### Hemodynamic Parameters of Interest

1. Fontan Efficiency: Indexed Power Loss (iPL)

iPL is a numerical representation of the power loss inside the Fontan conduit regarding each patient’s physiological conditions. The high iPL represents increased pressure changes in blood flow, which can exacerbate the cardiac performance (7). The iPL equation is as follows:

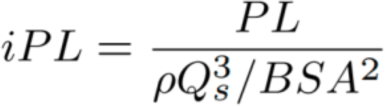

where ρ is the density, Q_s_ is the systemic venous flow, and BSA is the body surface area of the patient. The power loss PL is calculated by:

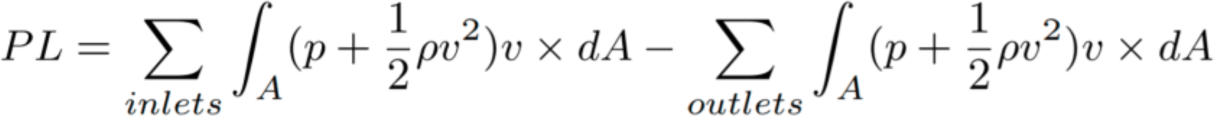

where A is the boundary area, *p* is the static pressure, and *v* is the velocity.

2. Hepatic Flow Distribution (HFD)

The HFD is the ratio of blood flowing to the pulmonary arteries (PA) from the inferior vena cava (IVC). The HFD was estimated through the Lagrangian particle tracking approach to track the blood flow streamlines in CFD. By releasing massless particles on the IVC boundary surface, the particles are traced to identify the number of particles that arrived at each ending of the PA (N_RP A_ and N_LPA_ respectively). HFD can thus be defined by the following equations:

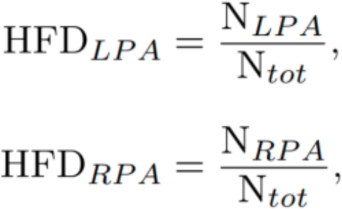

where N_tot_ represents the total number of particles that are released from the IVC.

### Boundary Conditions

Due to the passive venous flow from the cavae into the pulmonary arteries, prior studies have demonstrated accuracy of modeling continuous flow in the venous system (21). Previous simulation studies showed good agreement between using time-averaged and pulsatile BC in simulating Fontan hemodynamics (22). Therefore, in this study, the IVC and SVC flow rates were time averaged through summing up the phase velocity data, then divided by the overall time phase for inlet flow BC. The flow rates of the PA were assigned as the ratio of the flow split to the LPA and to RPA. The patient-specific data and the BC of the Fontan models are summarized in Table 1.

**Table 1:** Patient demographics (n=10)

| Case ID | Fontan Type | Cardiac Anatomy | BSA ($m^2$) | SVC (ml/s) | IVC (ml/s) | LPA (ml/s) | RPA (ml/s) |
| --- | --- | --- | --- | --- | --- | --- | --- |
| EX1 | Extracardiac | Hypoplastic Left Heart Syndrome | 1.457 | 21.69 | 43.21 | 23.57 | 41.33 |
| EX2 | Extracardiac | Hypoplastic Left Heart Syndrome | 1.593 | 23.05 | 40.29 | 27.55 | 35.78 |
| EX4 | Extracardiac | Dextrocardia-Unbalanced Atrioventricular Canal-Bilateral Glenn | 1.472 | L: 10.01, R: 17.22 | 58.91 | 42.21 | 41.9 |
| EX5 | Extracardiac | Dextrocardia-Unbalanced Atrioventricular Canal - Lglenn | 1.755 | 26.17 | 46.17 | 51.34 | 21 |
| LT1 | Lateral Tunnel | Levo-Transposition of the Great Arteries – remote ventricular septal defect | 1.984 | L: 5.62, R: 5.62 | 58.77 | 42.34 | 27.68 |
| LT2 | Lateral Tunnel | Double-inlet left ventricle – pulmonary atresia | 1.535 | 15.49 | 60.02 | 34.1 | 41.4 |
| LT4 | Lateral Tunnel | Tricuspid atresia | 1.645 | 23.46 | 48.1 | 42.87 | 28.69 |
| LT5 | Lateral Tunnel | Tricuspid atresia - Dextro-Transposition of the Great Arteries - Ventra Septal Defect | 1.625 | 16.21 | 32.91 | 23.68 | 25.45 |
| LT6 | Lateral Tunnel | Tricuspid atresia - bilateral SVC | 2.255 | L: 11.15, R: 10.84 | 71.2 | 60.45 | 32.83 |
| LT7 | Lateral Tunnel | Double-inlet left ventricle | 1.767 | 23.08 | 64 | 49.39 | 37.69 |

### *In Vitro* Validation Experimental Platform

#### 3D-Printing of Fontan Models

The *in vitro* study requires 3D-printing of Fontan models to be tested in the flow loop system that simulates the blood circulation in the total cavopulmonary connections. To connect the physical Fontan models to 15.875 mm (5/8”) tubing, a 30 mm extension was made on each inlet and outlet for fluid flow stabilization (Supplementary material Figure S2). To connect the eventual physical model to the tubing, a 20 mm long cylinder of the same diameter was added, separated from the previous extension by a loft of a determined length to create a 7 degrees wall angle to minimize the separation of the flow (23). A 3.175 mm (1/8”) shell was created around the model with open ends. This thickness is selected to match the thickness of the tubing the model would be connected to. The 3D models were printed by a third-party company (Xometry, Rockville, MD) using selective laser sintering (SLS) in Nylon 12 material.

#### Flow Loop Setup

*In vitro* validation of the simulation results was performed by setting up a flow loop to mimic the circulation of the 3D-printed Fontan models. Figure 2 shows the diagram of the system setup. 3D-printed Fontan models were connected to the experimental flow loop that consists of a flow pump (Flojet 04300143A Electric Pump, Xylem Inc), a fluid reservoir, Tygon tubing (OD 7/8”, ID 5/8”, McMaster-Carr), a customized compliance chamber, four inline ultrasonic flow meters (F-4600 Inline Flow Meter, ONICON inc.), and gate valves (2621-005G, PVC Valve, Spears Manufacturing Company). 60/40 water-glycerol mixtures were used to match the flow properties of blood (24–26).

**Figure 2.**
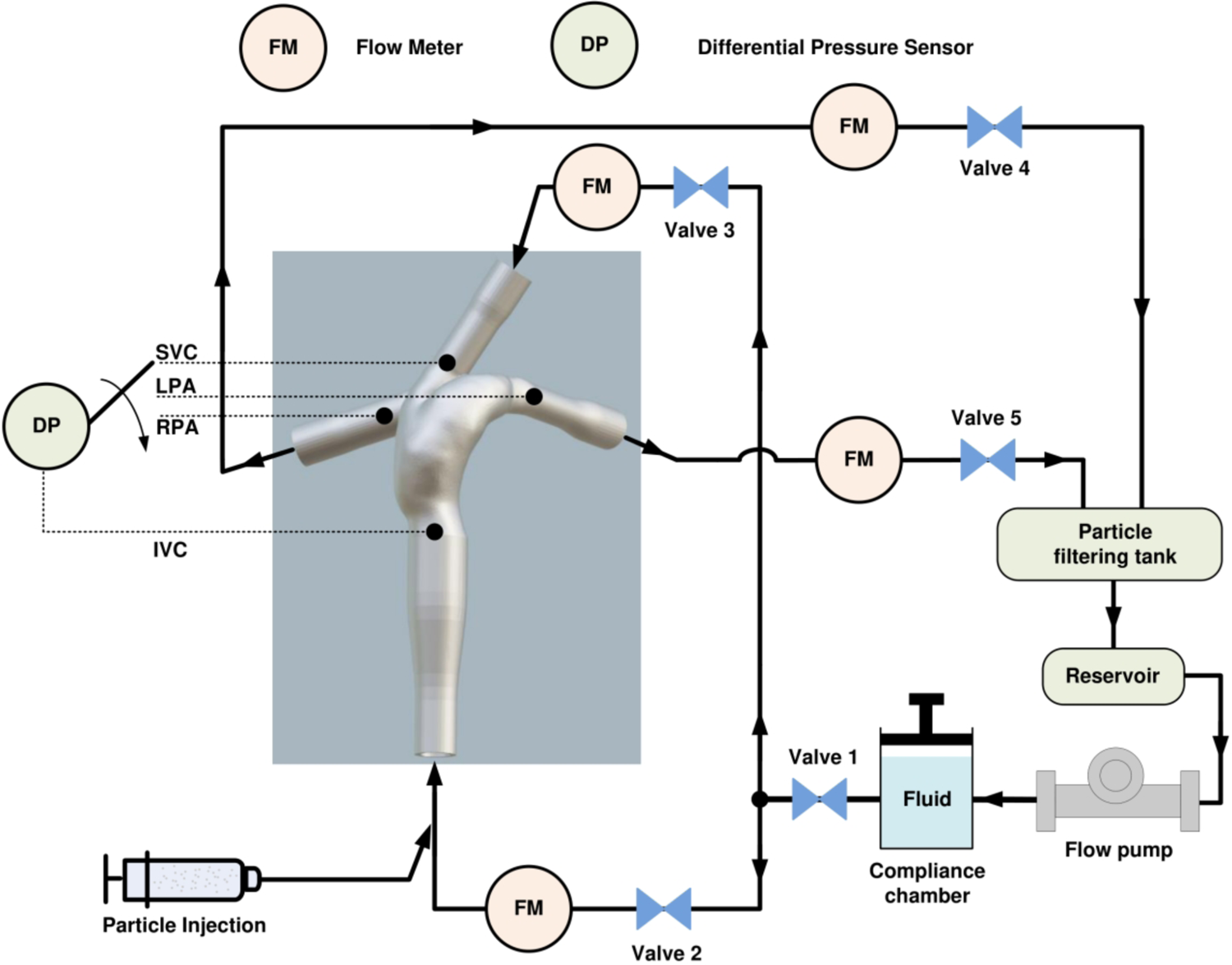
*In vitro* experimental setup. The experiment setup shows a 3D-printed Fontan model with two inlets and two outlets to connect with the flow loop. The fluid flow was generated by a flow pump with adjustable flow rates. Gate valves and flow meters (FMs) were used to adjust the flow rates that were prescribed at the inlets and outlets of the selected Fontan model. Microparticles were injected at the IVC and sorted by the particle filtering tank for measuring the hepatic flow distribution. A differential pressure sensor (DP) were used to measure the pressure differences of IVC-SVC, IVC-LPA and IVC-RPA for calculating the iPL.

#### Measurement of iPL

We employed a differential pressure sensor (Omega Engineering Inc.) with a resolution of 0.03 mmHg to measure the differential pressures (Supplementary material Figure S3). The PL can thus be calculated by using the measurements (Formula S1). To obtain robust and accurate measurements, we eliminate the effect of hydrostatic pressure (Δp_0_) from different poses of Fontan models by subtracting Δp_0_ from the measured Δp (Supplementary material Figure S4).

#### Measurement of HFD

Micro-polyethylene particles with diameters of 850-1000 µm (Cospheric LLC) were employed to suspend in water-glycerol mixture and inject to the tubing connected to IVC by using a syringe. The injected particles moved along with the fluid flow, exited from LPA and RPA, and were captured separately by a customized particle filtering tank (Supplementary material Figure S3). To make the microparticles well distributed in the fluid, we apply polysorbate as surfactant on the surface of the particles. The flow rate of particle injection from the syringe was controlled below 1 mm-s^−1^ to minimize the flow interruption at IVC. A customized particle dryer was used to dehydrate microparticles. An analytical scale with 0.001 g resolution was used to weigh the microparticles in different containers. The HFD is estimated by the weight ratio. (Supplementary material Figure S5)

### CFD Simulation

The workflow of the CFD simulation method is shown in Figure 3. The computational models were created by including extensions at the inlet and outlet boundaries of the models. The length of the extension at the inlet was chosen to be 10 times the inlet diameter as it produced insignificant differences in hemodynamics when compared with the results that were obtained by using real velocity profiles (27). The outlet boundaries were extended 50 mm to avoid backflow (9). The inlet extensions will significantly increase the computation time. To address the problem, the velocity field of each model using 1 mm mesh size with 10 times inlet extension was pre-computed. We extracted the velocity profiles 10 mm away from the actual SVC and IVC inlets (Supplementary material Figure S6).

**Figure 3.**
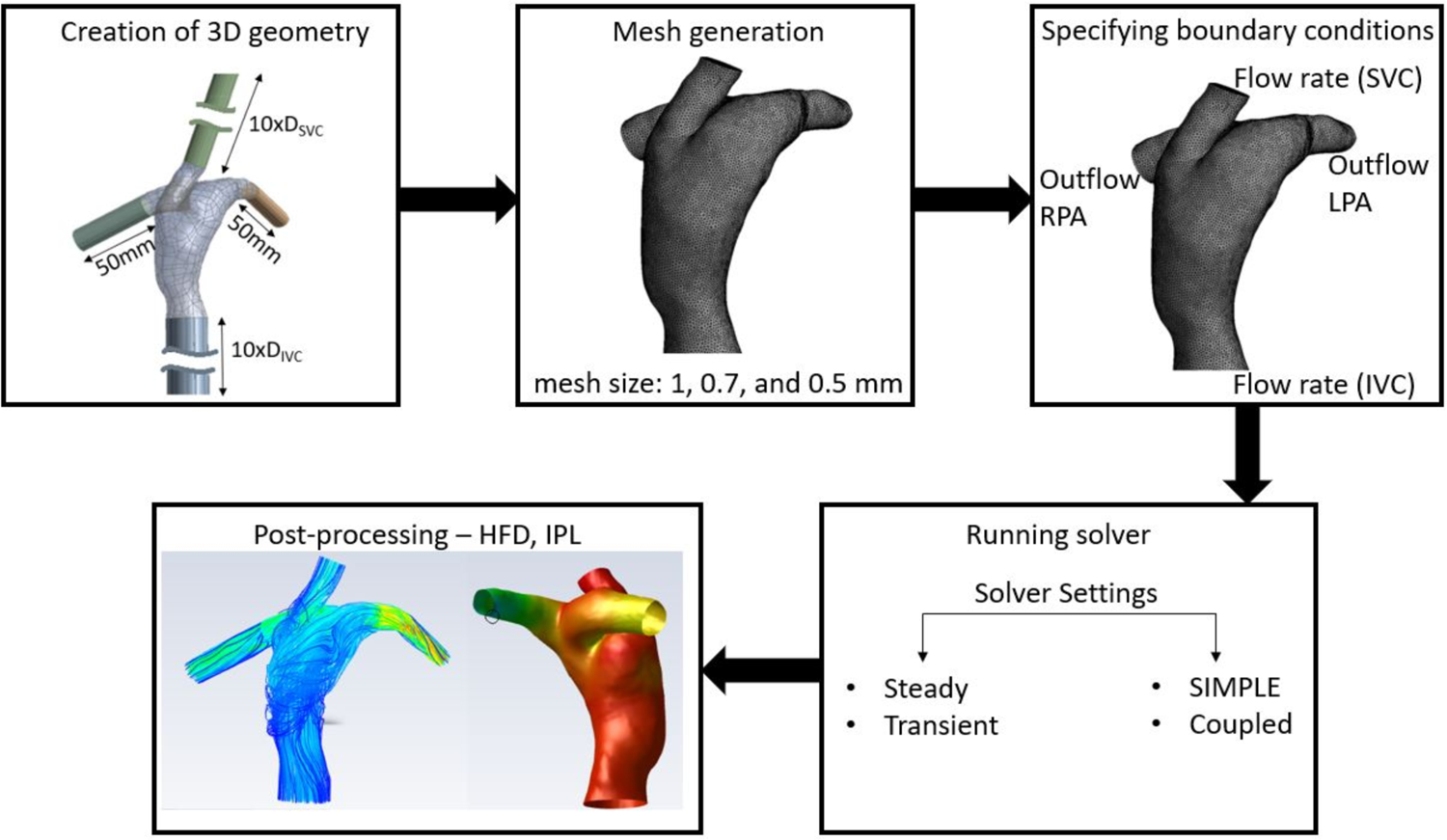
Procedure of CFD Simulation.

The total volume of each model was divided into small tetrahedral volumes with different mesh sizes 1.5, 1, 0.7, and 0.5 mm by using Ansys mesh. Seven-layer inflation with smooth growth was applied to the model walls, excluding extensions to better capture the velocity gradients.

Three types of solvers, including SIMPLE steady-state, coupled steady-state, and SIMPLE transient, were investigated by applying 0.5 mm, 0.7 mm and 1 mm mesh sizes to all models. Both the steady and transient flow equations formulated as follows were solved using time-averaged flow rates at the SVC and IVC inlets, and the percentages of total inflow at the LPA and RPA outlets (8). The BC of the Fontan models are presented in Table.

### Statistics

For all simulations and experiments, data are represented graphically as scatter plots of individual values with a bar identifying the median. Mean and standard deviation of each data group are represented as mean ± standard deviation. Comparisons of CFD and experiment results were analyzed using the paired 2-tailed t-test. A P value < .05 was considered statistically significant. Statistical analysis was performed by using Prism version 9 (GraphPad Software, California).

## Results

### Comparison of CFD and *In Vitro* Results

Because of the significant impact of different mesh sizes and the insignificant impact of different solvers on the CFD results (Supplementary material Figure S7 and Figure S8), we compared the CFD results with the experimental results by using 0.7 mm and 1 mm by using the SIMPLE-steady solver, as shown in Figure 4A and Figure 4B. The green, blue, and red bars indicate the CFD results using 1) 0.7 mm, 2) 1 mm mesh sizes, and 3) the experiment results for iPL and HFD, respectively.

**Figure 4.**
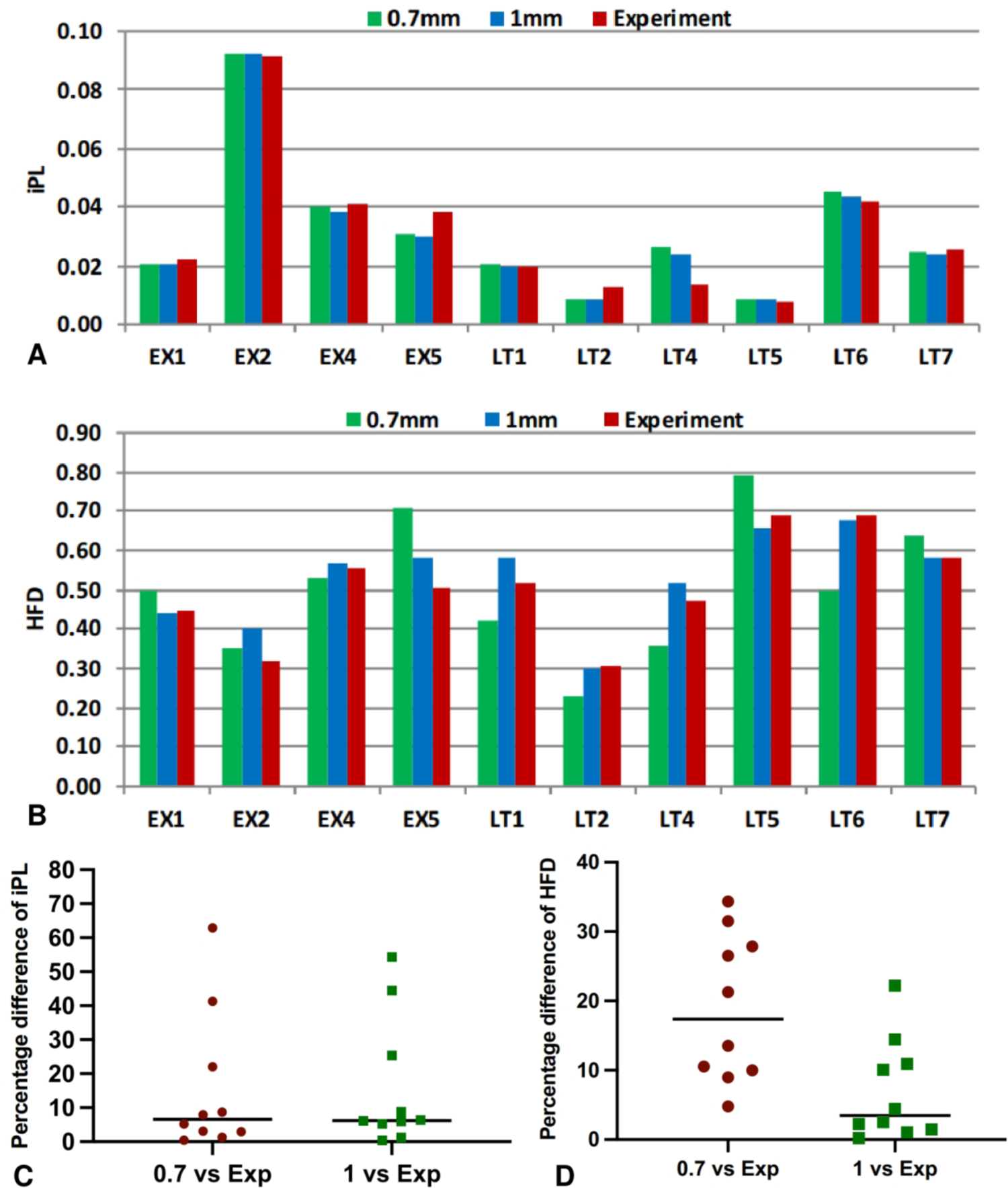
Comparison of iPL and HFD between CFD and experimental results. The CFD results were computed by using 0.7 mm and 1 mm mesh sizes separately with the SIMPLE-Steady solver.

Figure 4C and Figure 4D show the percentage difference of iPL, HFD with two comparison groups: 1) 0.7 mm mesh CFD simulation vs experimental results (0.7 vs Exp), and 2) 1 mm mesh CFD simulation vs experimental results (1 vs Exp). Figure 4C shows the medians of the percentage difference of iPL between CFD and experiment are smaller than 10% for “0.7 vs Exp” (15.57%±20.83%) and “1 vs Exp” (15.82%±19.12%). The two groups have no significant difference P = 0.9775. In contrast, Figure 4D shows the HFD simulation results with 1 mm mesh size have a significantly smaller difference with the experimental results than those of 0.7 mm mesh size (P = 0.0087). The medians, means, and standard deviations are 17.39%, 18.93%±10.63% for “0.7 vs Exp”; and 3.47%, 6.95%±7.25% for “1 vs Exp”.

### Computation Efficiency

The computation time depends on the computing power, number of iterations, mesh sizes, and numerical solver types. As the mesh size decreases, the number of mesh elements increases. Using different solvers (steady SIMPLE, steady coupled, transient SIMPLE) and various mesh sizes (0.5 mm ∼ 2.5 mm) for the simulations, the required computation time for 3000 iterations (24 CPU cores for each simulation) was shown in Figure 6 for one of the models (EX5). This is a representation of how the time that requires to complete the simulations varies by using different solvers and mesh sizes. The steady-SIMPLE solver with the 2.5 mm mesh size shows the highest computation efficiency (9.2 minutes) for providing over 90% iPL and HFD prediction accuracy. While the transient solver with 0.5 mm mesh size shows the lowest computation efficiency (937 minutes). Despite of the fast computation speed by using 2.5 mm mesh size, we tend to keep the mesh size below 1 mm for preventing potential simulation failures with the excessively large meshes.

**Figure 5.**
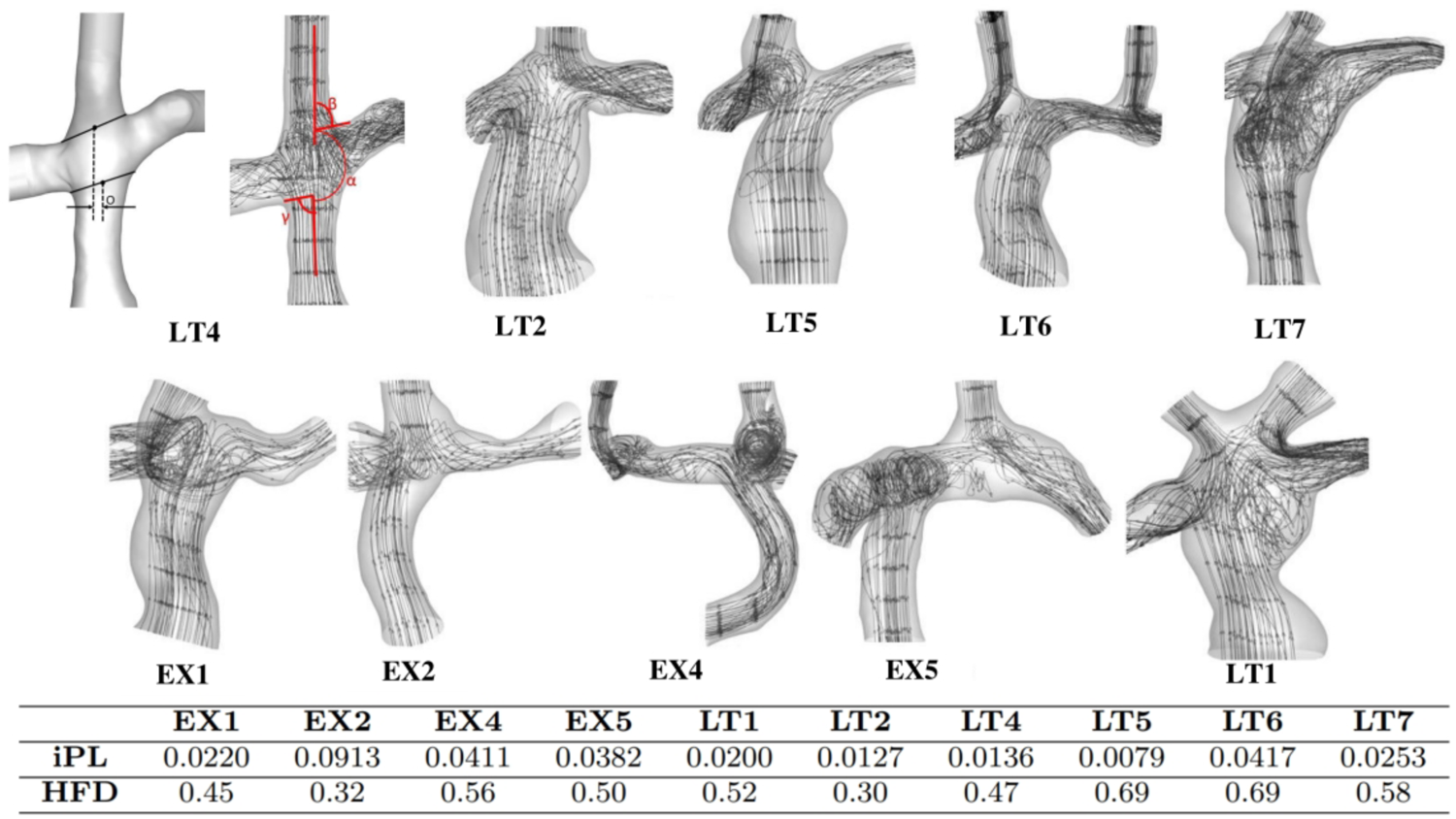
Analysis of Fontan model geometry and flow streamlines. The offset (O) measured between the center of IVC and SVC where the flows enter the PA, angle between the IVC and SVC flow where they enter the PA. The table at the bottom shows the angle and offset between the flows from IVC and SVC.

**Figure 6.**
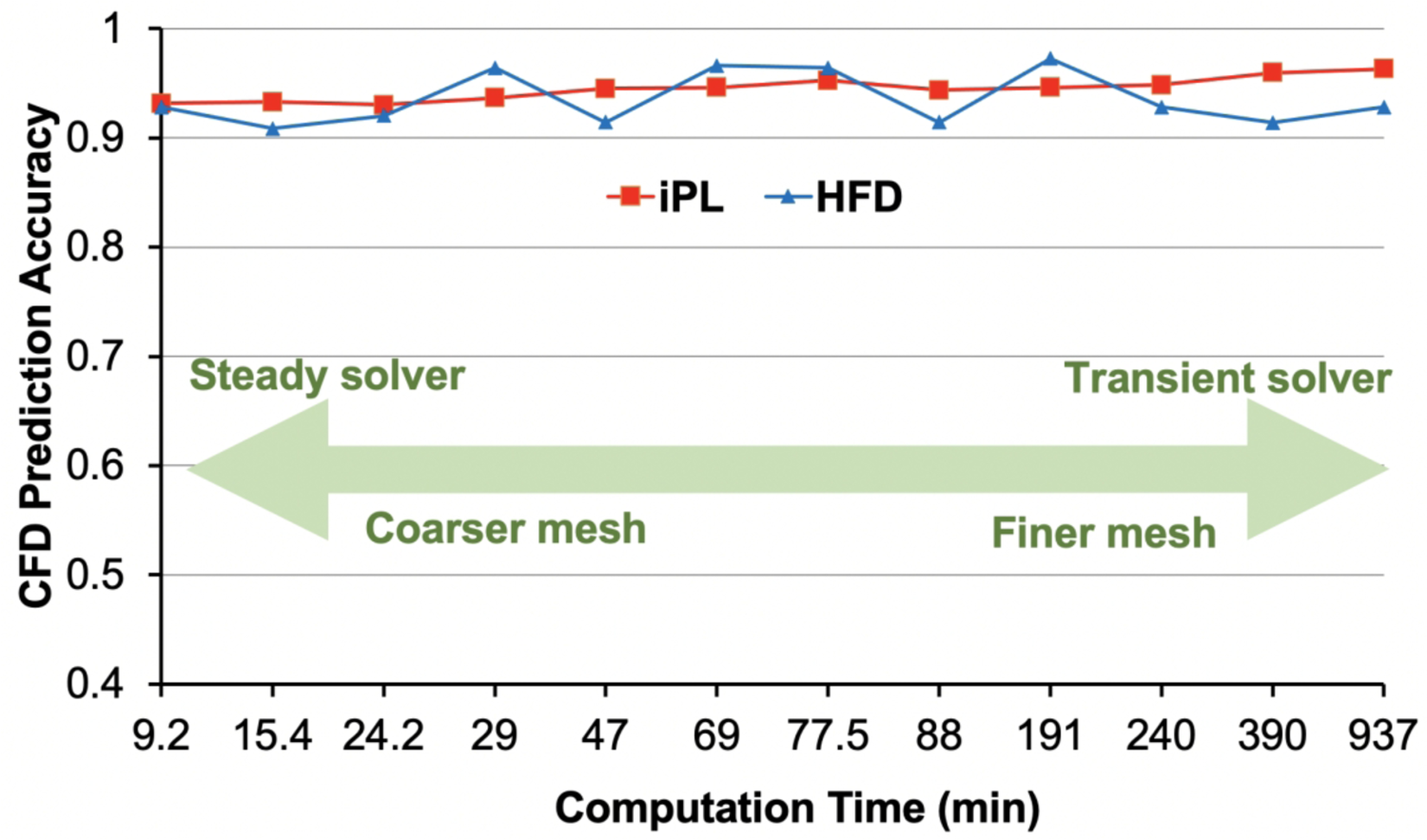
Computation time and CFD prediction accuracy by using different solvers and various mesh sizes to complete the simulations with 3000 iterations on EX5.

### Discrepancy Analysis for iPL

Large discrepancy of iPL between the experimental and CFD results was seen in models: LT2, LT4, and EX5. To investigate the sources of this discrepancy, the anatomy of the models and flow fields were compared. First, the offset between the IVC and SVC centerline and the angle between the SVC and IVC flow entering the PA were presented in Figure 5. The Fontan model LT4 was used to show where the angle and offset were measured. The flow streamlines for the models were shown in each model. The flow competition between IVC and SVC flows is evident, especially for the models with smaller offsets and larger angle. The models, which have higher discrepancy in iPL (EX5, LT2, and LT4), have the IVC and SVC flows entering the PA with almost 90° (γ and β, respectively). The vortices are formed and the flow in the PA becomes more complex and difficult to model when the inflows hit the PA wall or each other at 90°. Furthermore, the region where the SVC and IVC flows hit each other is where most of the power is dissipated. For example, 78% and 64% of the total power in LT2 and LT4 is dissipated in this region, respectively.

## Discussion

Fontan surgical planning by using CFD analysis has gained a great attention from clinicians to provide valuable insight into possible post-surgical outcomes and reduce uncertainty in decision making and personalizing surgical approaches for each patient. However, CFD research has not yet translated into broad clinical acceptance because most surgical planning processes are primarily operated by engineering teams, and the surgical planning has thus been relegated to retrospective post-hoc analysis. In clinical application, the required time between pre-operative imaging that is used for CFD analysis and Fontan surgery can vary, but on average takes around two months (3, 28). Considering that at least 1000 Fontan operations occur nationally (29), computational resources will start to become an issue. In this study, we found that there are diminishing returns on investment for both mesh size and solver sophistication of CFD simulation; and can recommend standards to provide clinically acceptable Fontan predictions, allowing for realistic clinical use. We also identified that there are certain Fontan geometries that may be prone to more discrepancies and would warrant clinicians to be cautious of the CFD results of these types of Fontan geometries.

This study aimed to quantify both the accuracy and efficiency of different CFD setups for Fontan surgical planning, validated by a sophisticated *in vitro* bench setup. A key innovation of this work was development of an *in-vitro* micro particle tracking system for reliable HFD *in vitro* measurements. Prior research studies employ 4D flow MRI data and particle tracking method to estimate HFD (16–18). However, the 4D flow MRI itself may require validation by using PIV (20) or CFD simulation (21) due to its limited spatial resolution, signal noise, and the segmentation errors. To the best of our knowledge, this is the first use of an *in vitro* flow loop setup to validate HFD with a large enough cohort size for a statistical comparison.

The clinical impact of iPL is with regards to the patient’s exercise capacity that is known to deteriorate over time (30), while the HFD, which is diversion of normal hepatic venous flow from the pulmonary circulation, is related to the development of PAVM (31). In our CFD mesh analysis, we found iPL is more independent on mesh size than HFD. There were insignificant differences in iPL by using 0.5 mm∼1.5 mm mesh sizes. In addition, there were insignificant differences by using different numerical solvers for both iPL and HFD. Therefore, if a study only focuses on iPL, the 1 mm mesh size with the SIMPLE steady solver can be a good starting selection for CFD simulation because of 1) the small standard deviation (1.42%) of the percentage difference of iPL by comparing with the results from the 0.5 mm mesh size, and 2) the fastest computation speed of the SIMPLE steady solver.

Different from iPL, there is a significant difference in HFD between the mesh sizes above 1 mm and below 0.7 mm. We found that the HFD results computed with 1 mm match significantly closer to the experimental results than those computed with 0.7 mm. The Fontan models with the 0.7 mm mesh size have denser massless particles uniformly distributed at the inlet boundary surfaces than those with the 1 mm mesh. In our *in vitro* experiment, we used microparticles with the size of 0.85 mm∼1 mm. The distribution of the microparticles released at IVC in the experiment matches closer to the simulated particle distribution with the 1 mm mesh in the simulation than that with the 0.7 mm mesh size.

Based on our results, the cumulative probability distributions of the percentage difference between the CFD predicted values and the experimental benchmark values for iPL and HFD are generated (Supplementary material Figure S9). For a CFD simulation of a patient-specific Fontan model, the probabilities of keeping the percentage differences of iPL and HFD below 10% are about 0.6 and 0.8, respectively. Our analysis indicates that the flow competition of IVC and SVC can lead to a significant power loss, which agrees with the finding in a prior study (32). The computational modeling with different solvers used in this study seems insufficient to accurately compute the power loss under this scenario. To predict iPL accurately for these complex models, further computational analysis is needed.

While robust and novel, our work was limited by the limited size variety of the microparticles to fully confirm the influence of mesh size on HFD, as they are not readily commercially available. In addition, we acknowledge that a larger scale of validation is still needed to generate more robust statistics on the accuracy of CFD prediction for cardiovascular surgical planning, which include, but not limited to Fontan surgery. Surgical implantations of planned vascular grafts were executed by using a paper printed ruler as guidance to match the prescribed anastomosis location (13). More work is required to quantitatively analyze and improve the implantation accuracy. To robustly acquire the flow measurements, during the experiment, we oriented the 3D printed Fontan models in different poses to mimic different orthostatic positions and adjusted the fluid density and viscosity by changing the percentages of water-glycerol mixture from 60/40 to 80/20. Our results show that the measurement changes of iPL and HFD are small enough to ignore. The Nylon polymer used in the 3D printed Fontan model is more rigid than the actual Gore-Tex graft that can resist dilation. Rigid wall assumption is appropriate to be used in Fontan analysis. Prior fluid-structure interaction study showed minimal differences in HFD by comparing rigid and compliant vessel wall simulations (33). To avoid geometric confliction of Fontan conduit and heart, at the phase of surgical planning, the heart model was used to define the spatial constraint. Further clinical study is needed to confirm how graft-heart interaction affects Fontan hemodynamics. Beyond the scope of this work, effects of non-compliant vessel/chamber walls, respiratory flow changes, flow changes with exercise and the patient’s growth, anatomical changes with the patient’s growth will be investigated in future study.

## Conclusions

Our results show that there are 0.6 to 0.7 probability chances for the CFD simulation to predict the iPL and HFD with higher than 90% accuracy, which indicates that CFD simulation can provide meaningful hemodynamic prediction for Fontan surgical planning. The prediction accuracy can be significantly affected by the model shape complexity and flow competition.

## Supplementary Documents

A summary of the supplementary materials is demonstrated in the file Supplementary Materials.docx with a list of figures provided separately.

## Acknowledgements

This work is supported by the National Institutes of Health under award numbers R01HL143468 and R21/R33HD090671. The authors acknowledge the supercomputing resources at the University of Maryland (http://hpcc.umd.edu) and the Maryland Advanced Research Computing Center (MARCC) (https://www.marcc.jhu.edu) that made available for conducting the research reported in this paper.

## Glossary of Abbreviations

3D: Three-dimensional

BC: Boundary Conditions

CAD: Computer-Aided Design

CFD: Computational Fluid Dynamics

HFD: Hepatic Flow Distribution

iPL: Indexed Power Loss

IVC: Inferior Vena Cava

MRA: Magnetic Resonance Angiography

LPA: Left Pulmonary Artery

PA: Pulmonary Arteries

PAVM: Pulmonary Arteriovenous Malformations

PL: Power Loss

RPA: Right Pulmonary Artery

SCPC: Superior Cavopulmonary Connection

SVC: Superior Vena Cava

SIMPLE: Semi Implicit Method for Pressure Linked Equations

---

**Figure S1:**
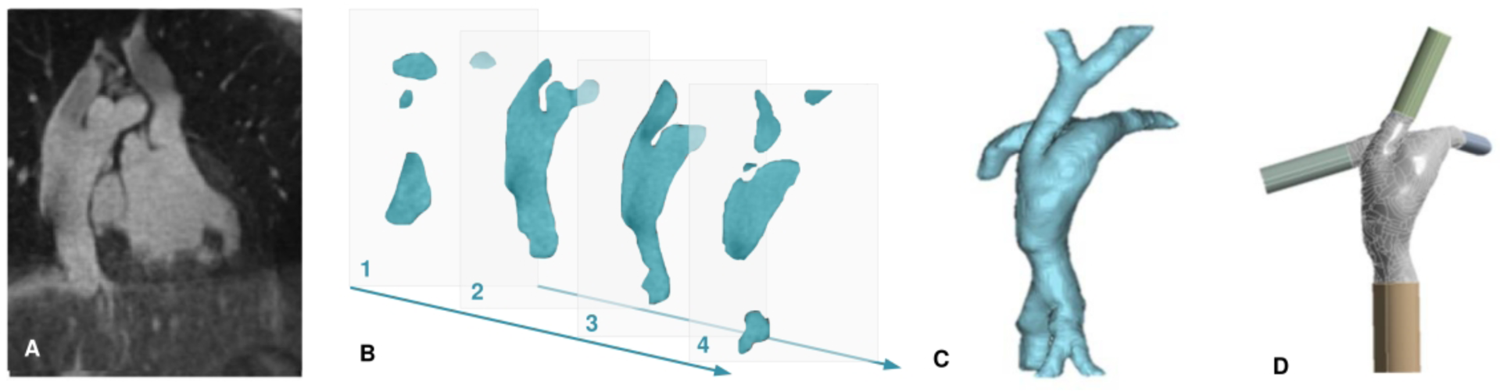
Schematic workflow of patient-specific Fontan model preparation. (A) Data acquisition by contrast-enhanced magnetic resonance angiography (MRA). (B) The MRA dataset slices with the region of interest, the Fontan blood pool. (C) 3D Fontan model constructed by the MRA slices. (D) Preparation of the model with extensions at the boundaries for CFD simulation.

**Figure S2:**
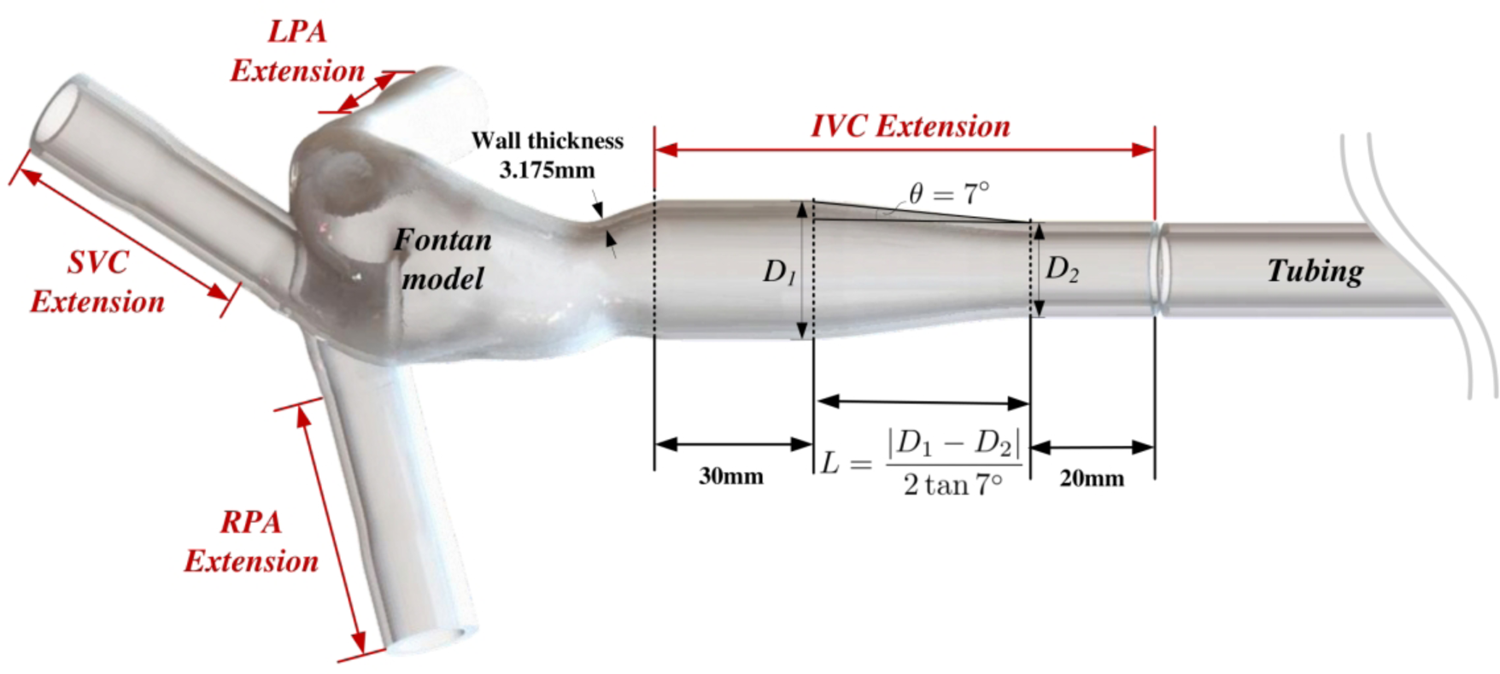
Illustration of the inlet/outlet extensions for the 3D printed Fontan models.

**Figure S3:**
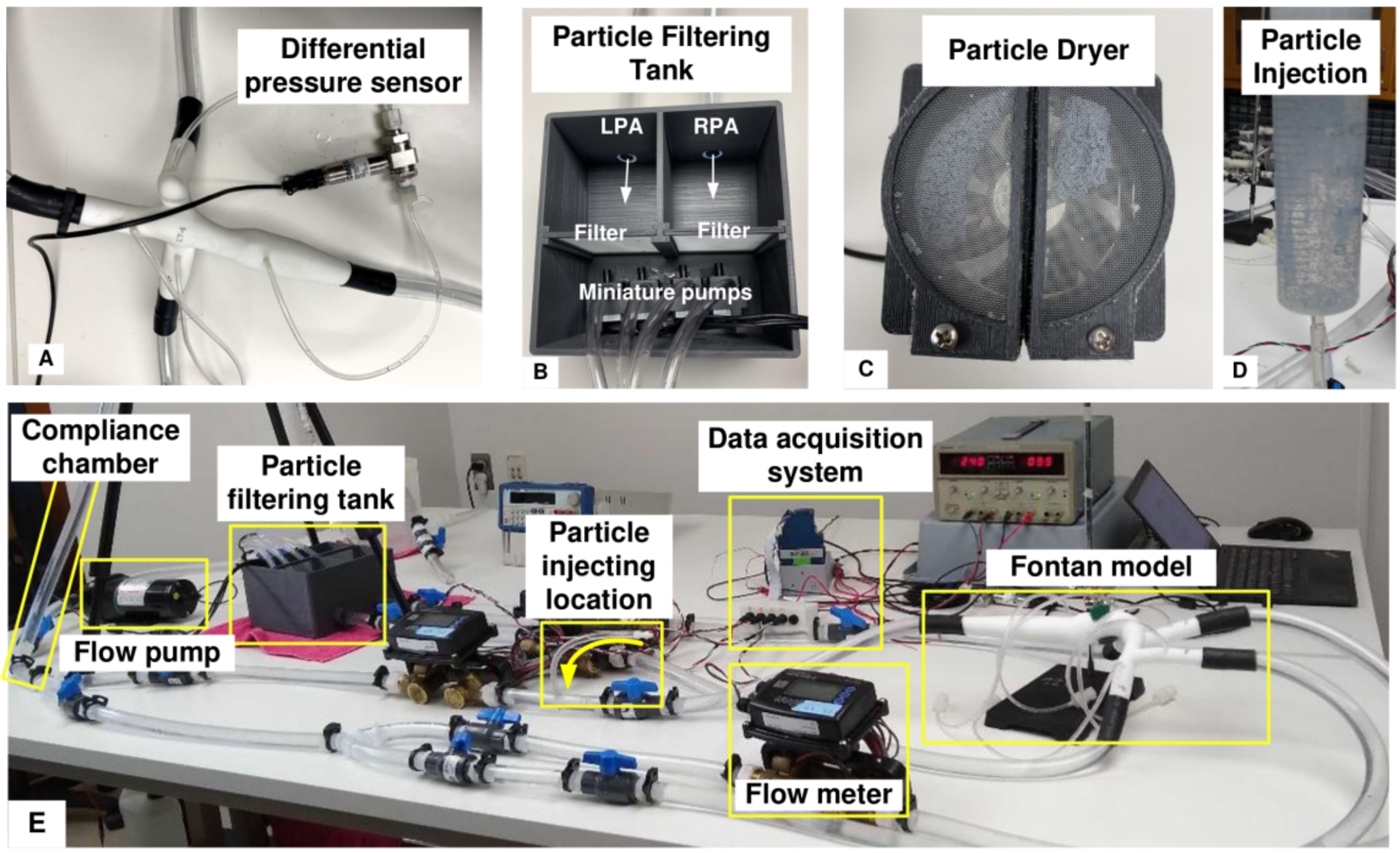
Physical experiment setup. (A) A 3D-printed Fontan model with DP attached. (B) The particle filtering tank with two separated containers to sort the microparticles exited from LPA and RPA. The miniature pumps were used to prevent liquid building up in the tank. (C) The particle dryer. A cooling fan installed at the bottom was used to dehydrate the particles for accurate weight measurement. (D) The syringe loaded with fluid with suspended microparticles. (E) The overview of the physical experimental platform.

**Figure S4:**
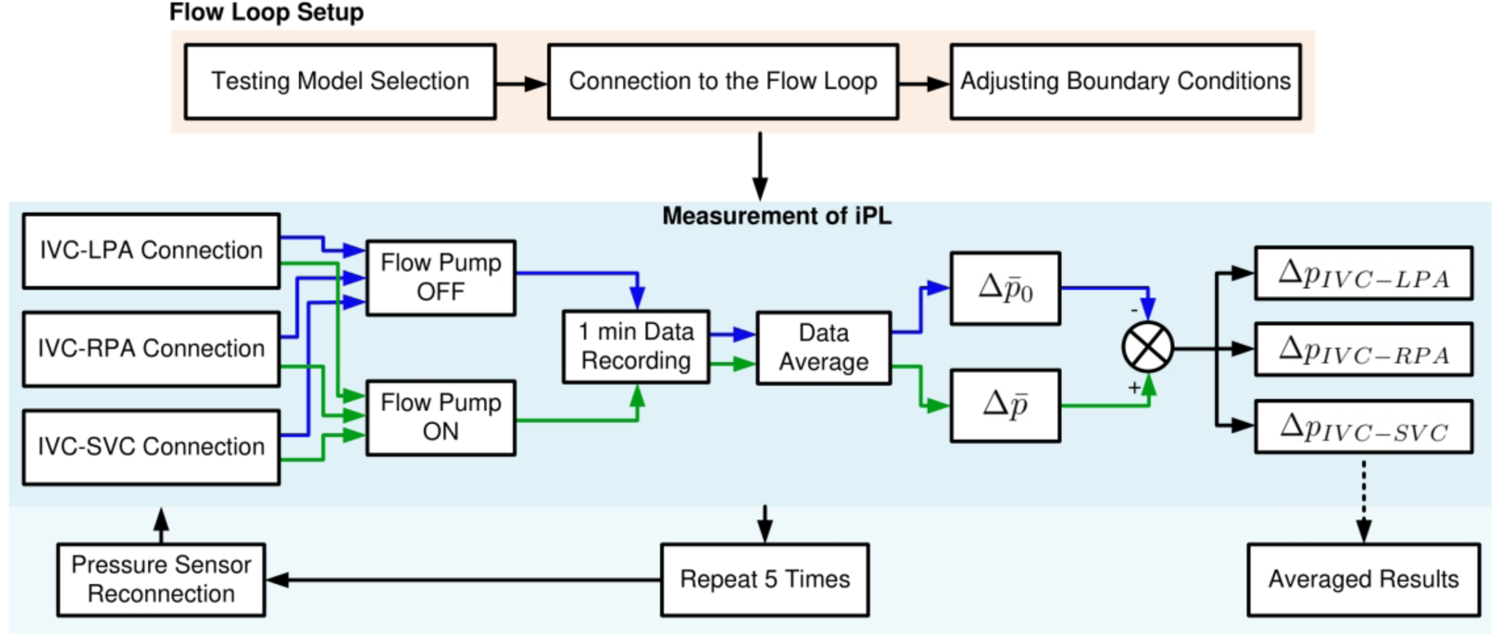
Experiment procedure of measuring iPL. By setting up the Fontan model and adjusting the BC, the differential pressure sensor (DP) was connected to IVC-LPA, IVC-RPA, IVC-SVC separately. For each connection, the differential pressures were recorded by turning the flow pump ON and OFF for cancelling out the influence of model orientation related hydrostatic pressure.

**Figure S5:**
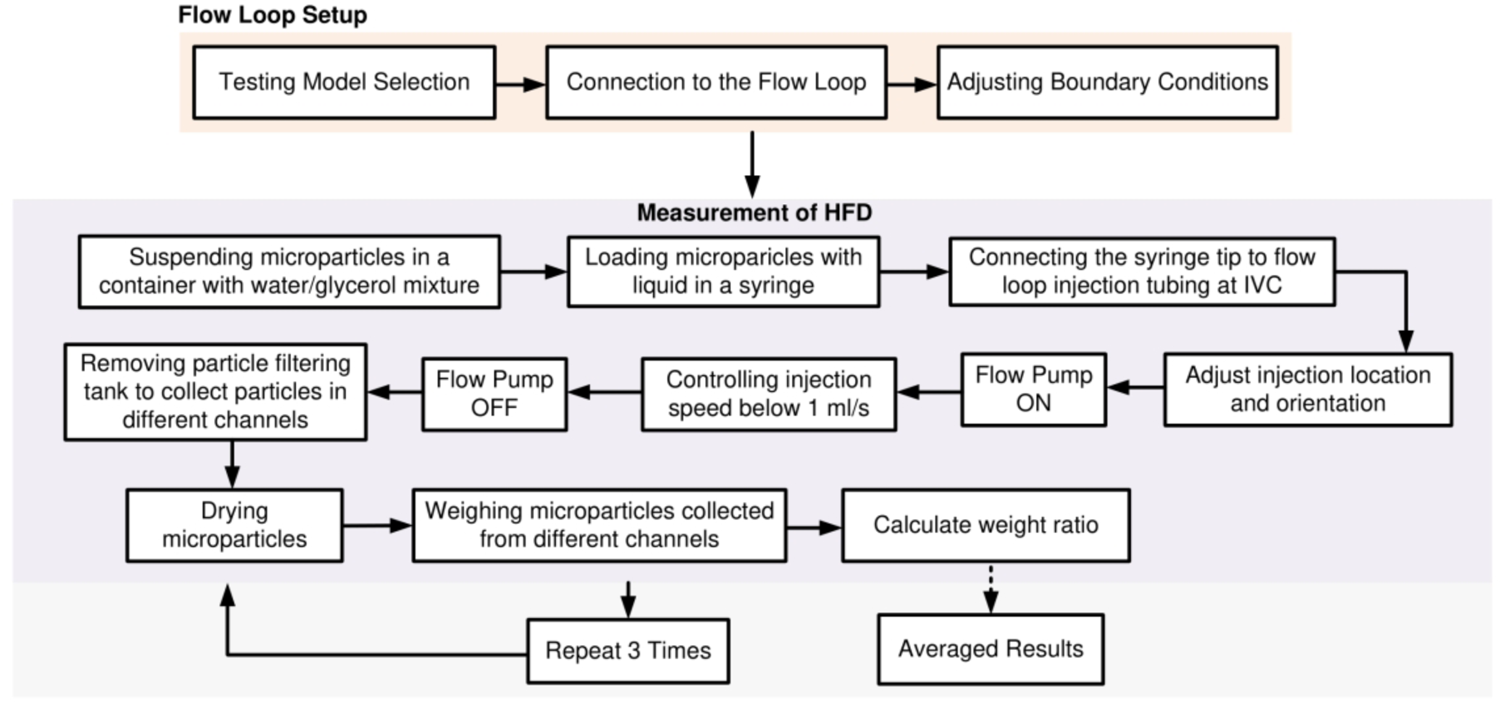
Experiment procedure of measuring HFD. The microparticles were prepared to suspend in the liquid by applying surfactant. A syringe was used to load the liquid with the microparticles and to inject at the IVC with the injection speed less than 1 mL/s. The micropaticles were captured by a particle filtering tank and dehydrated by a particle dryer. The weights of microparticles collected from LPA and RPA were measured separately by using an analytical scale.

**Figure S6:**
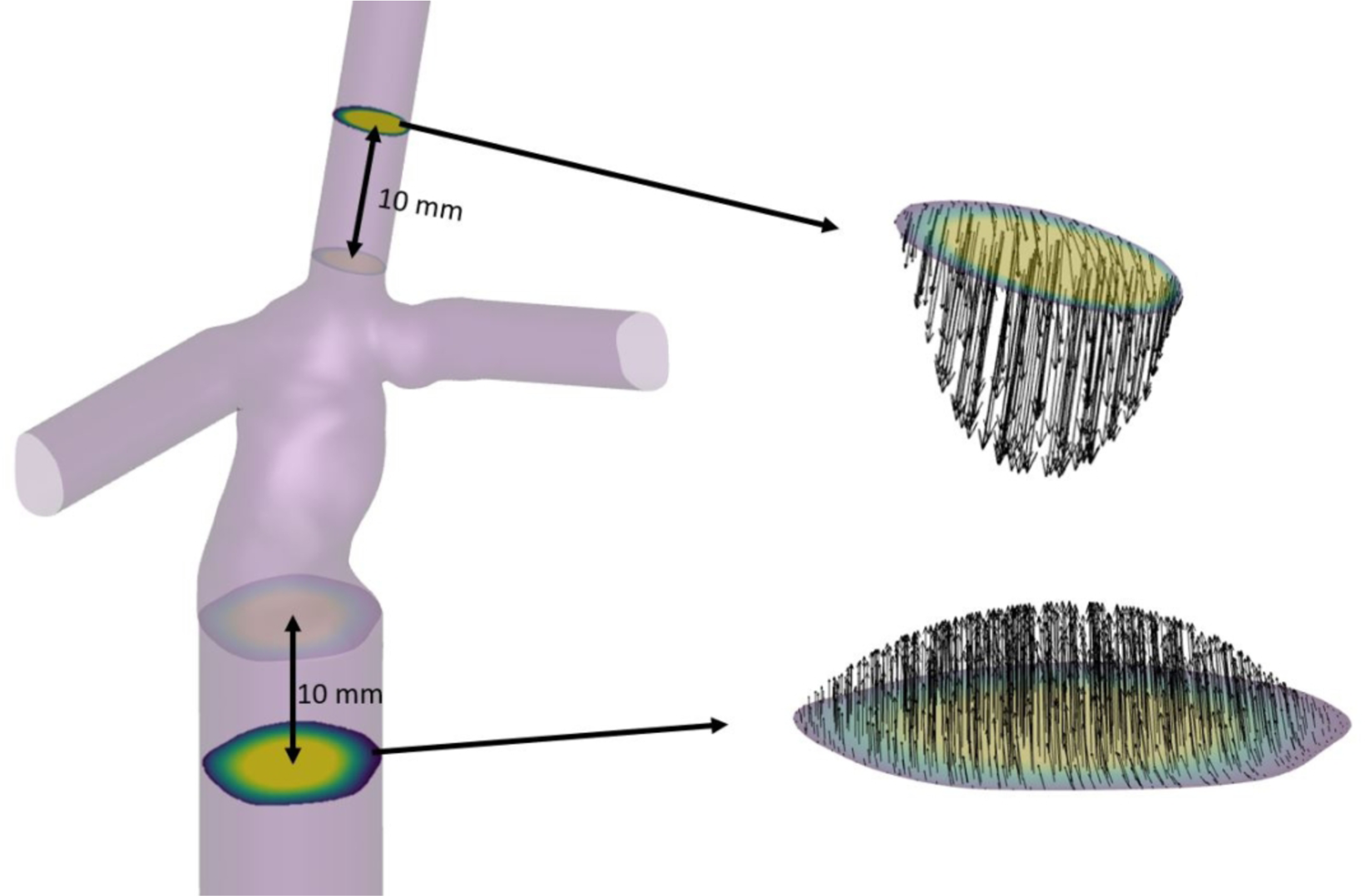
Illustration of velocity profile extraction at the inlets.

**Figure S7:**
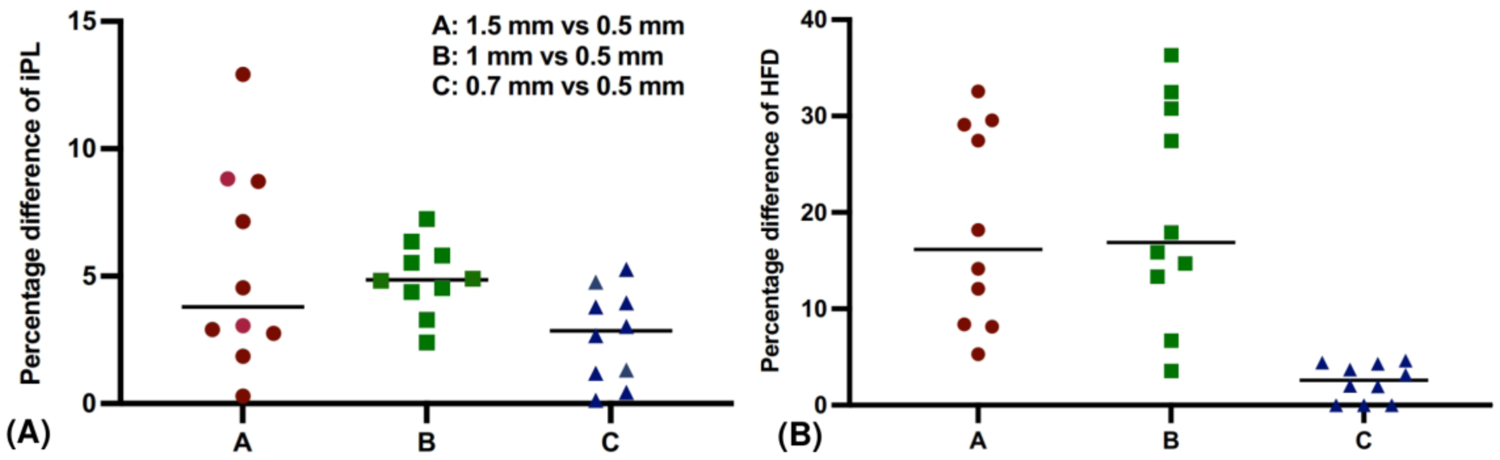
Mesh analysis for computing iPL and HFD by using the SIMPLE steady solver. To demonstrate the mesh convergence, the iPL and HFD results were obtained by using 0.5, 1, 0.7, and 1.5 mm mesh sizes. We compared the percentage differences of iPL and HFD results between the 0.5 mm mesh size and the other mesh sizes for each model. Three comparison groups including 1.5 vs 0.5 (short for comparison between 1.5 mm mesh size and 0.5 mm mesh size, and so on), 1 vs 0.5, 0.7 vs 0.5 were used for this analysis. There is no significant difference among 1.5 vs 0.5, 1 vs 0.5, and 0.7 vs 0.5 for the percentage difference of iPL, P=0.0697. (A) 0.5vs 1.5, 5.30%±3.95%; 0.5 vs 1, 4.93%±1.42%; 0.5 vs 0.7, 2.66%±1.82%. In contrast, there is a significant difference among the three groups for the percentage difference of HFD P = 0.0002. (B) 0.5 vs 1.5, 18.50%±10.31%; 0.5 vs 1, 19.92%±11.24%; 0.5 vs 0.7, 2.43%±1.91%.

**Figure S8:**
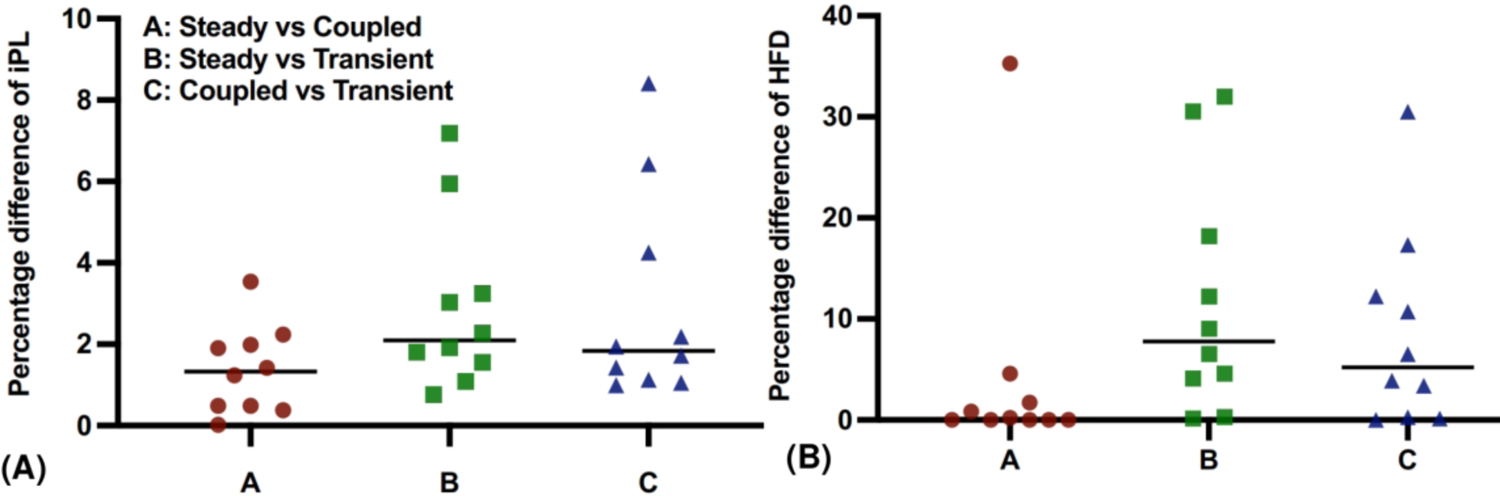
Numerical solver analysis for computing iPL and HFD by using 0.7 mm mesh size. According to the mesh analysis results, we employed 0.7 mm mesh size to investigate how different solvers affect the iPL and HFD results. Similar to the mesh analysis, we calculated and compared the percentage difference of iPL and HFD among the SIMPLE-steady, coupled, and SIMPLE-transient solvers. (A) The percent differences in iPL were less than 10%. There is no significant difference between the solvers for iPL (steady vs coupled, 1.37%±1.08%; steady vs transient, 2.88%±2.11%; coupled vs transient, 2.96%±2.58%; P = 0.5017). (B) The percentage differences in HFD, especially for groups B and C, were larger than those in the iPL. There is no significant difference between the solvers for HFD (steady vs coupled, 4.26%±11.00%; steady vs transient, 11.76%±11.62%; coupled vs transient, 8.50%±9.67%; P = 0.5541).

**Figure S9:**
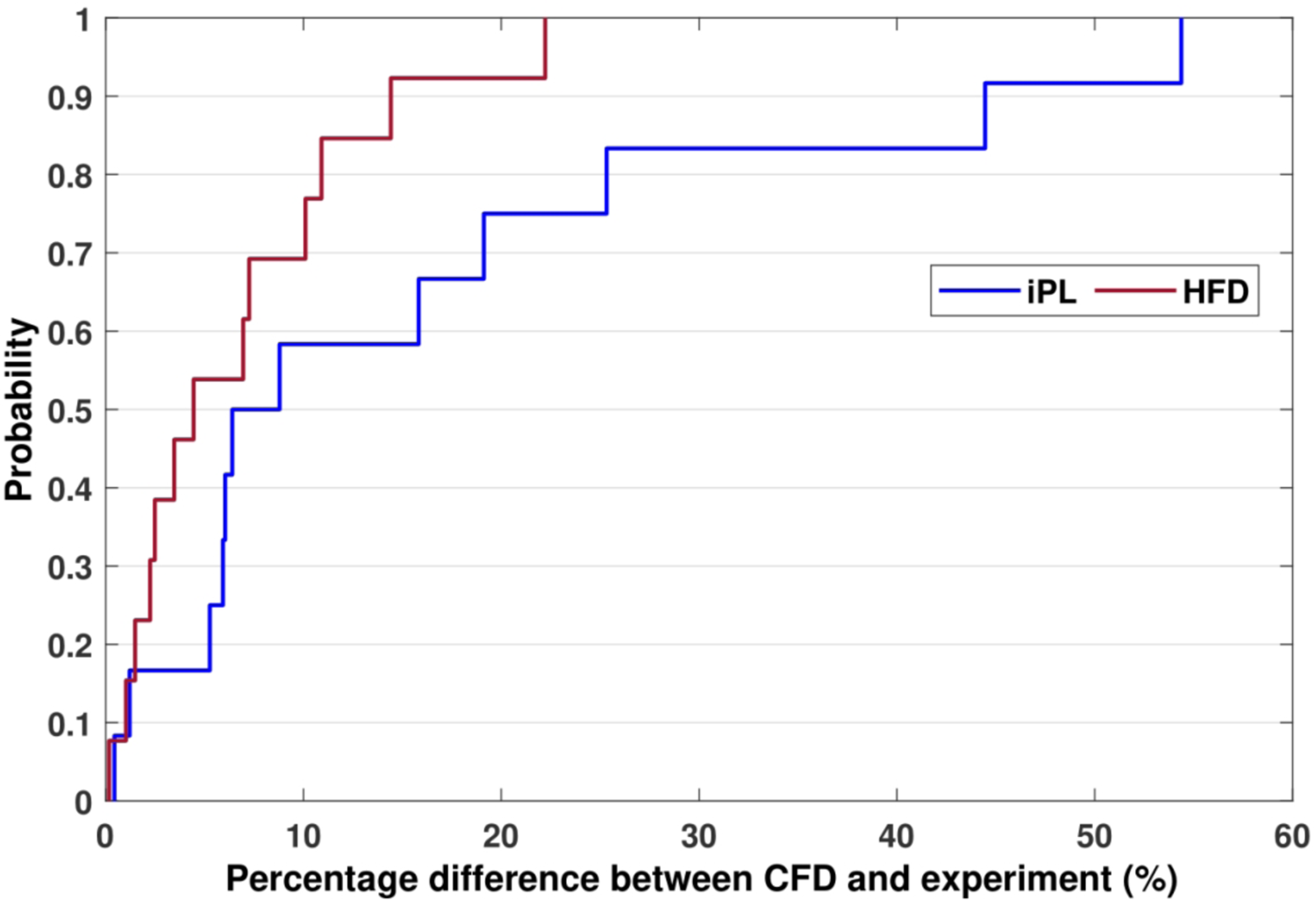
Cumulative probability distributions of the percentage differences between the CFD and the experiment results for iPL and HFD.

## Formula S1: Power Loss Calculation by Using Differential Pressures

The differential pressures are represented as

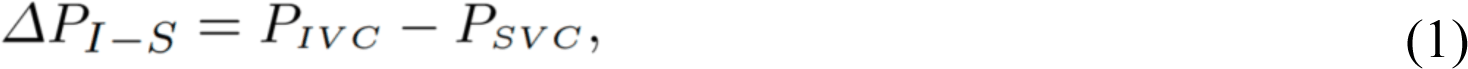

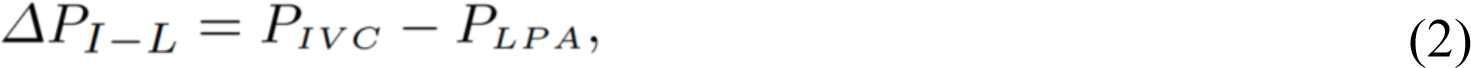

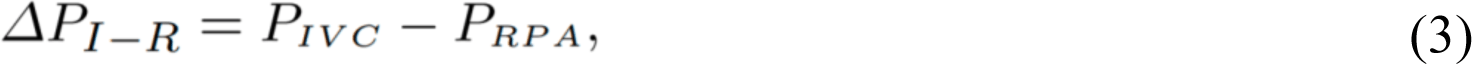

where *P_IVC_*, *P_SVC_*, *P_LPA_* and *P_RPA_* are absolute static pressures at the inlets and outlets, ΔP_I−S_, ΔP_I−L_, and ΔP_I−R_ are the differential pressures that can be acquired by the differential pressure sensor. From (1), (2), (3), the absolute static pressures are denoted as

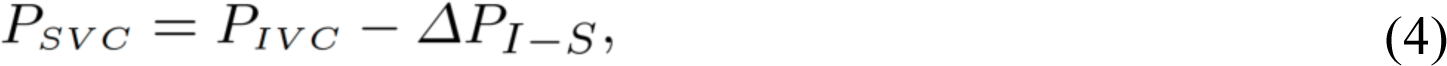

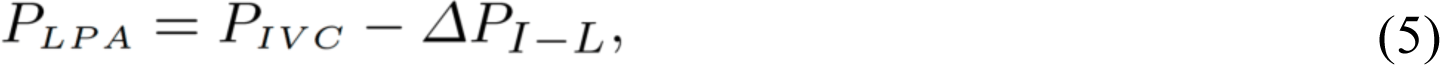

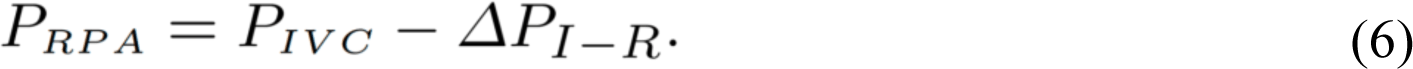

By expanding (7),

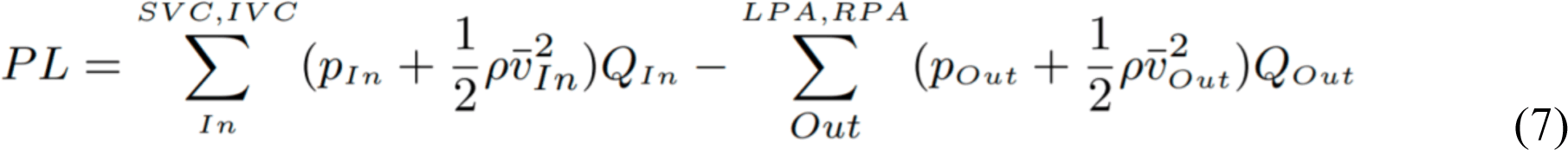

we have

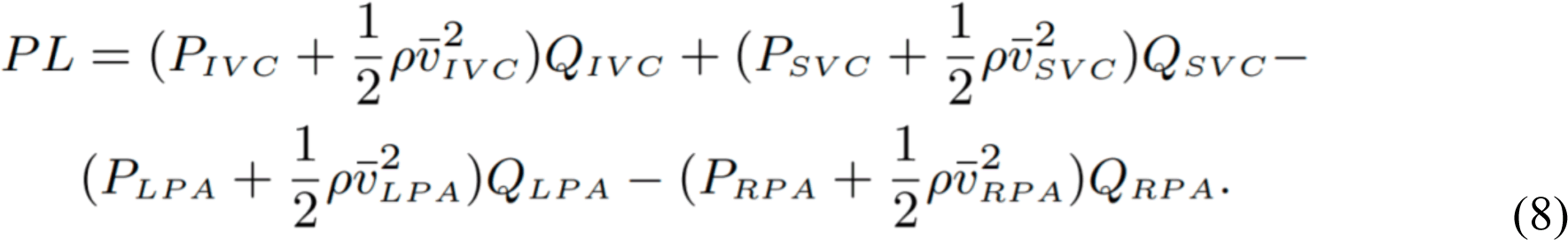

By substituting (4), (5) and (6) in (8), we have

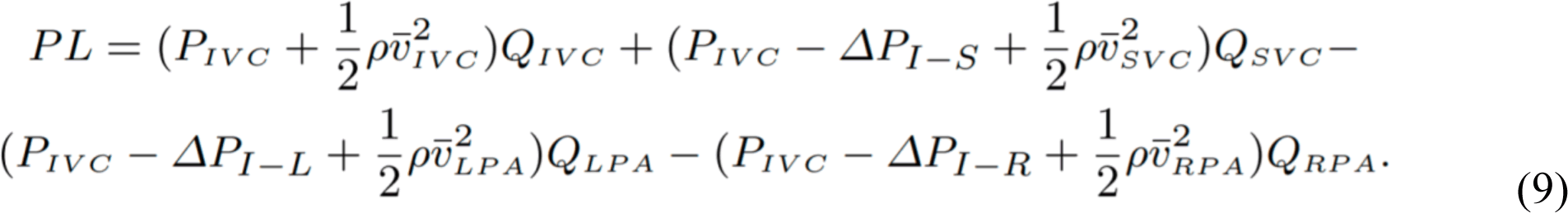

By rearranging (9), we have

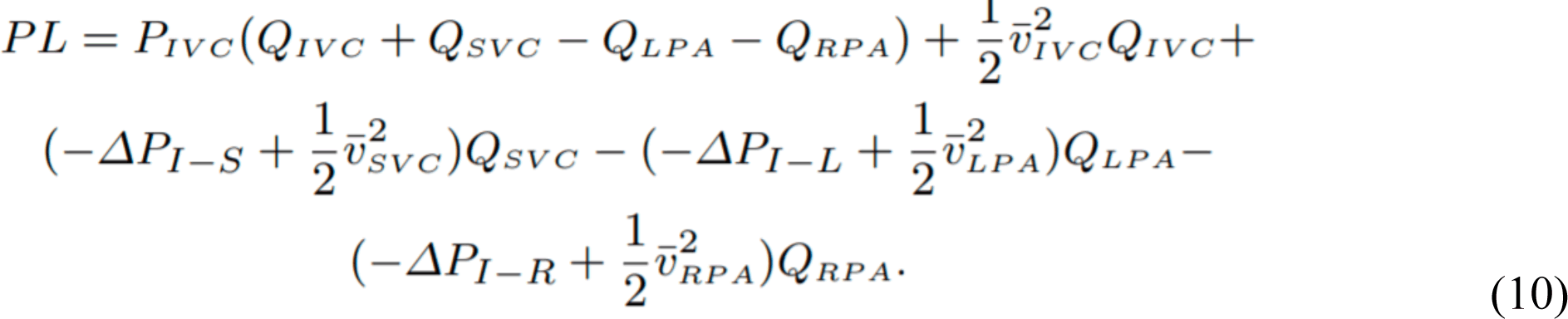

According to the conservation of mass flow rates at the inlets and outlets, we have

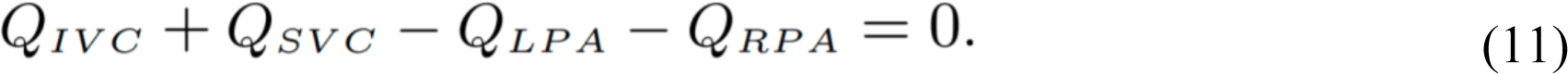

Therefore, the first term in (10) can be cancelled out. By rearranging (10), we can have the PL equation based on differential pressures

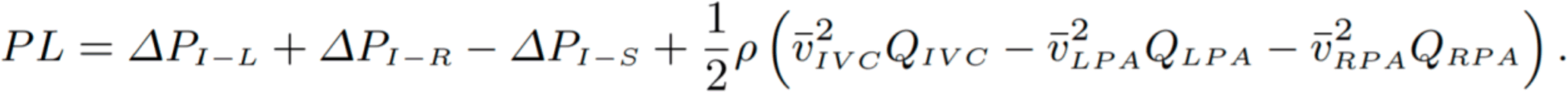

